# Analysis of chronic host-aspergilloma interactions using a novel mouse model

**DOI:** 10.1101/2023.10.06.561242

**Authors:** Ryosuke Hamashima, Masato Tashiro, Yuichiro Nakano, Hotaka Namie, Yuya Ito, Tatsuro Hirayama, Kazuaki Takeda, Naoki Iwanaga, Kodai Nishi, Hong Liu, Takahiro Takazono, Takeshi Tanaka, Akira Watanabe, Yoshihiro Komohara, Akitsugu Furumoto, Katsunori Yanagihara, Hiroshi Mukae, Scott G Filler, Koichi Takayama, Koichi Izumikawa

## Abstract

An aspergilloma is a fungus ball caused by chronic infection of *Aspergillus* species in a pre- existing cavity, such as a destroyed lung or the sinuses. Patients with pulmonary aspergilloma are at risk of sudden life-threatening hemoptysis. Antifungal therapy is administered to aspergilloma patients who are ineligible for surgery, but its efficacy is limited. Understanding the pathophysiology of aspergilloma is crucial for developing further treatment strategies. The mechanism behind the long-term host response to aspergilloma is poorly understood. We created a novel mouse model to analyze the host response to aspergilloma by implanting a fungus ball of *Aspergillus fumigatus* into an air-filled subcutaneous cavity. Our findings indicate that a live fungus ball led to tissue invasion, even in immunized mice. When a fungus ball consisting of dead hyphae was implanted, it persisted for over three months and induced pathological findings simulating human aspergilloma, including an inflammatory cell infiltration into the fungus ball and angiogenesis in the cavity wall. Dead fungus ball induced Th1 and Th17 inflammatory cytokines and vascular endothelial growth factor. Neutrophils infiltrated the inside of the fungus ball immediately after implantation, and macrophages surrounded it after a one-week delay. The macrophages around the fungus ball were swollen with phagocytosed fragments of dead hyphae and transformed into foam cells containing fat droplets. We also confirmed in vitro that macrophages were damaged and transformed into foam cells by direct contact with dead hyphae. This model holds promise to provide new insights into the fungal-host interaction during aspergillomas.

**NOTATION OF PRIOR ABSTRACT PUBLICATION/PRESENTATION:** None

**IMPORTANCE:** Chronic aspergillosis, which affects over 3 million people worldwide with a 5-year survival rate of 50%, is understudied compared to other forms. Our study focuses on aspergilloma, a key aspect of chronic aspergillosis, using a groundbreaking mouse model that mirrors clinical features over three months. By studying host-fungal interactions within aspergillomas, we discovered Th1 and Th17 inflammatory responses to dead fungal hyphae. Initially, neutrophils dominate, later giving way to macrophages with a lipid-accumulating foamy phenotype. This transition may impede aspergilloma clearance. In addition, even dead hyphae induce vascular endothelial growth factor and promote angiogenesis. Our findings, which are critical for the prevention of fatal hemoptysis, highlight the need for innovative treatments that target fungal clearance and challenge the limited efficacy of antifungal agents against dead fungal bodies. This research represents a significant step forward in the understanding of chronic aspergillosis.

## INTRODUCTION

An aspergilloma is a fungus ball that forms in preexisting air-filled spaces, typically lung cavities or sinuses, due to prolonged infection by *Aspergillus* species [1]. Lung cavities often result from chronic pulmonary diseases, such as tuberculosis, non-tuberculous mycobacteria, and chronic obstructive pulmonary disease that result in structural damage to the lung [2]. Pulmonary aspergillomas can cause symptoms such as hemoptysis and purulent sputum, while sinus aspergillomas can lead to facial pain and purulent nasal discharge [3,4]. In patients with pulmonary aspergilloma, approximately 23-30% experience massive hemoptysis, which can be life-threatening [5,6]. Additionally, approximately 22% of patients with pulmonary cavities after pulmonary tuberculosis will develop aspergillomas [7], leading to chronic pulmonary aspergillosis (CPA). There are an estimated 3 million cases of CPA worldwide, further emphasizing the global impact of this disease [8]. Therefore, due to the substantial risk and the high prevalence of CPA, aspergilloma is a clinically significant infection requiring attention and appropriate management strategies.

Aspergillomas typically needs surgery for effective treatment because antifungals have limited efficacy [9]. The 10-year survival rate of patients with aspergillomas is 84.8% when surgery can be performed, but drops to 56.7% when surgery cannot be performed [3]. Because the lungs of many people with aspergilloma have substantial damage, nearly 50% of these patients cannot undergo surgery [3]. Hence, there is a demand for crafting innovative treatment approaches for patients with aspergillomas.

A thorough comprehension of the pathogenesis of aspergillomas is necessary for developing innovative approaches to treatment. Aspergillomas are composed of fungal hyphae, inflammatory cells, fibrin, mucus, and cellular debris [10]. Most of the fungal hyphae in aspergillomas are dead, with only the surface hyphae remaining viable [11]. Clinical studies have demonstrated decreased staining within the interior of aspergillomas in surgical specimens [11], as well as low rates of positive cultures from sinus fungus balls (16.7-25.7%) [12–14].

There is typically no evidence of tissue invasion in the histopathological findings of aspergillomas as compared to invasive aspergillosis. The cavity containing an aspergilloma often has a disrupted epithelial lining due to prior lung diseases such as chronic obstructive pulmonary disease and old tuberculosis [15]. A study on the pathology of CPA reported that erosions are present in the cavity surrounding the fungus ball in all cases and constitute an average of 62% of the cavity tissue [15]. Chronic inflammatory changes in the cavity lining are variable, often neutrophilic with occasional eosinophilic exudates [15]. Notably, the rich capillary bed of granulation tissue in the cavity wall is generally the source of hemorrhage.

While there has been substantial research into how the immune system responds to invasive aspergillosis and allergic bronchopulmonary aspergillosis, almost no studies have focused on the immune response to aspergillomas. One significant challenge has been the absence of appropriate animal models for investigating aspergillomas [16,17]. Recently, several animal models that simulate *Aspergillus* colony formation in the airway have been reported [18,19].

These studies involved injecting conidia embedded in agar beads or small hyphal balls of less than 250 nm into the airways, with an observation period limited to 28 days [18,19]. However, this duration falls short of the minimum of at least three months that is necessary for studying clinical aspergilloma [9]. Creating an effective aspergilloma animal models is difficult due to the complexities of inducing cavities in small animals and the prolonged experimental timeline required by the chronic nature of the disease. Previously reported procedures for generating aspergillomas in animal models have been invasive and complex, such as ligating rabbits’ bronchi and blood vessels to induce bronchiectasis or creating a space in the chest cavity through artificial pneumothorax [20–22]. To overcome these issues, we attempted to make the cavity formation process less invasive and simpler by utilizing the subcutaneous space. To address the question of whether the cavities should be subcutaneous, we hypothesize the following: First, aspergilloma is not a tissue-specific disease, as it can develop not only in lung cavities but also in the sinuses and, rarely, in the middle ear after tympanic membrane perforation [1,23]. Secondly, in the pathology of pulmonary aspergilloma, the cavity wall is often eroded and fibrotic, and the original airway epithelium is lost [15]. Based on these clinical observations, although the airway is an important route of entry for *Aspergillus* spores, we believe that the presence of a “cavity” is the most important factor for the development of aspergillomas. Based on this hypothesis, we have successfully created a mouse model in which a fungus ball is introduced into an air-filled subcutaneous cavity on the back. Our mouse model provides important insights into the long-term host response and sheds light on the underlying reasons for the failure of the host immune system to eradicate aspergillomas.

## RESULTS

### A live fungus ball invaded the tissue even in healthy and immunized hosts

The sagittal cross-sectional image of our mouse model obtained from a Computed Tomography scan is shown in Figure 1. We found that live fungus balls can invade tissues, even in healthy mice (Fig. S1A and S1B). Many patients with aspergillomas become seropositive for *Aspergillus* antibodies after prolonged exposure to the organism [24]. Therefore, we sought to immunize the mice by administering solutions containing homogenized hyphal components of the fungal ball. We observed that 66.7% of the mice were positive for *Aspergillus* precipitins at four weeks after the start of immunization, with a subsequent increase to 100% at six weeks (Fig. S2). Even in these immunized mice, tissue invasion occurred upon implantation of live fungus balls (Fig. S1C, S1D), indicating that immunization does not prevent tissue invasion. Regardless of the immunization status, the majority of mice implanted with live fungal balls eventually experienced skin or spinal cord invasion, resulting in skin lesions or spinal cord injury within two weeks of implantation, precluding further observations.

**Figure 1.**
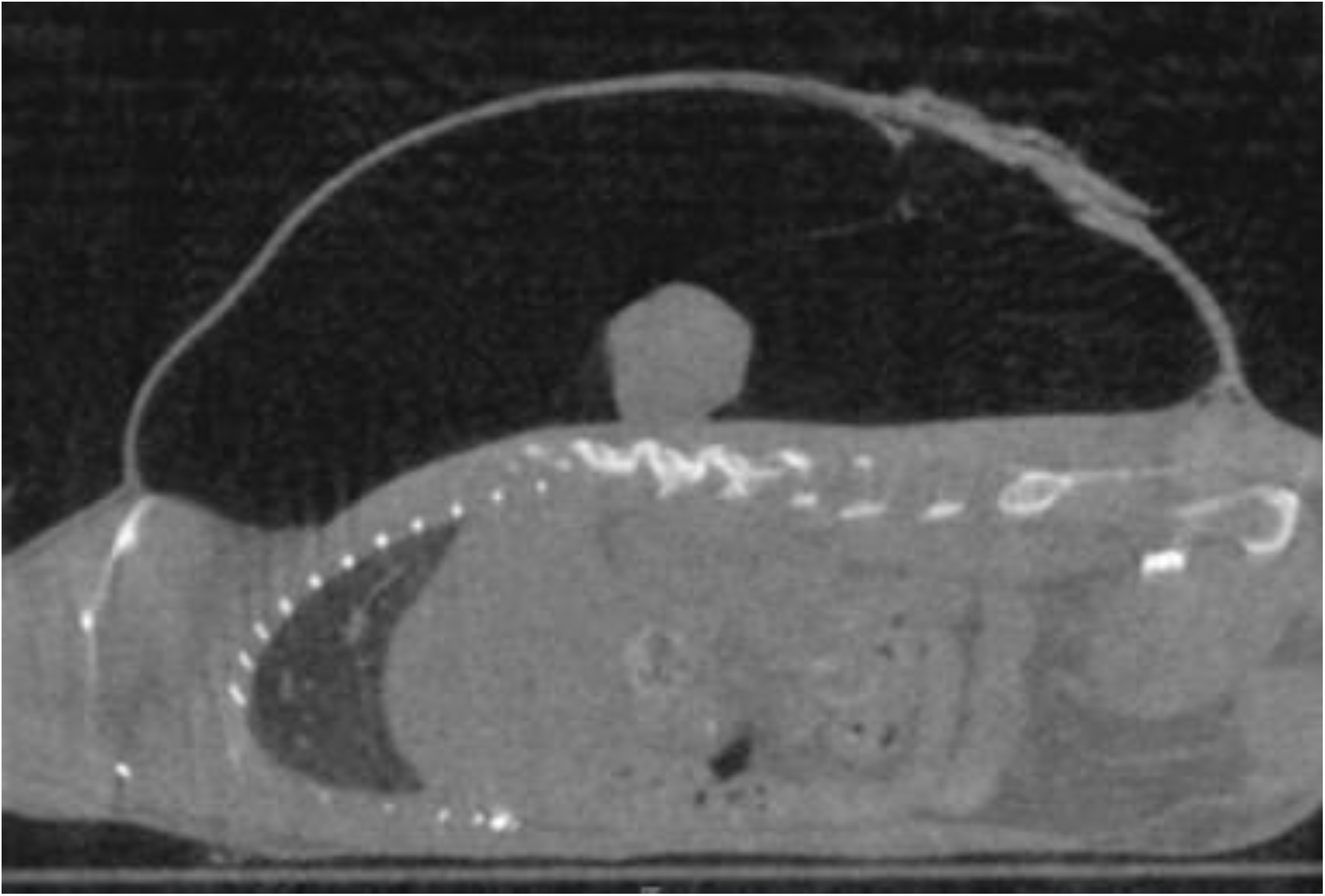
The sagittal cross-sectional image of our mouse model. The fungus ball is located on the muscle layer.

### The histopathology of mice with dead fungus balls is consistent with clinical aspergilloma

To avoid tissue invasion, we implanted dead fungus balls into healthy mice. The implantation of heat-killed fungus balls allowed for prolonged observation, in contrast to the implantation of live fungus balls. On day 104, histopathologic analysis of a mouse bearing a killed fungal ball revealed its persistence in the subcutaneous cavity (Fig. 2A, 2B). There was neutrophil and macrophage infiltration on the surface of the fungus ball (Fig. 2C). Notably, the surrounding wall of the fungus ball showed fibrosis and angiogenesis (Fig. 2D). These findings were consistent with clinical findings in that there was marked inflammatory cell infiltration of the fungus ball and fibrosis and angiogenesis in the cavity wall.

**Figure 2.**
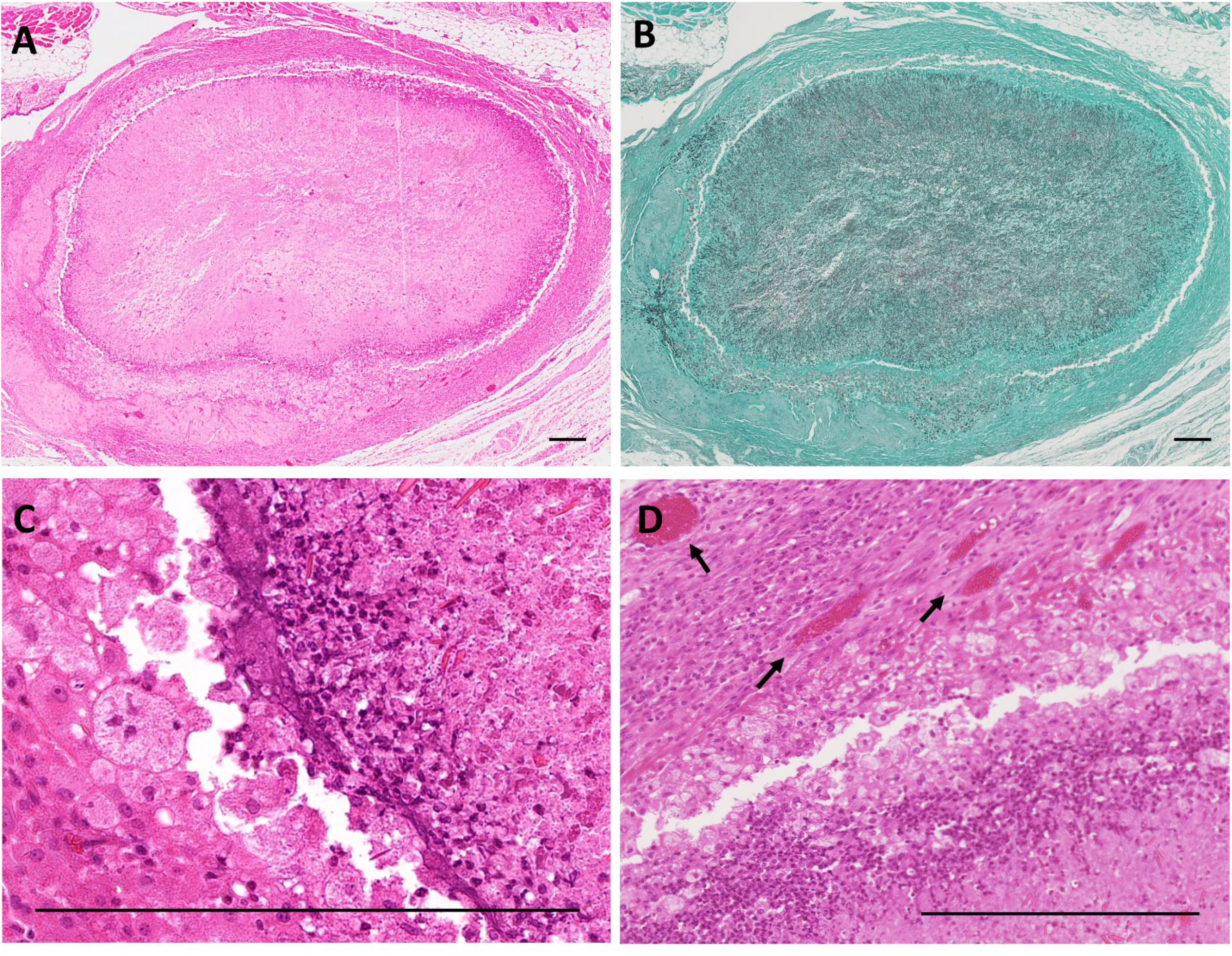
Histopathology of a mouse on 104 days after heat-killed fungus ball implantation. H&E staining (A) and GMS staining (B) are shown. Inflammatory cell infiltration inside and on the surface of the fungus ball (C) and fibrosis of the cavity wall (D) are seen. Black arrows indicate angiogenesis. Bars, 250 μm.

### Changes of dead fungus balls over time

The heat-killed fungus balls had a whitish appearance until day 7. By day 14, they assumed a yellowish-brown color with purulent characteristics (Fig. 3A). However, on detailed examination at days 104 and 161, the fungus ball became enveloped by a membranous structure and assimilated into the host tissue. Histopathological analysis demonstrated the presence of inflammatory cell infiltration inside the fungus ball as early as the day after implantation, while fibrous tissue encapsulation was observed surrounding the fungus balls after day 104 (Fig. 3A). To quantitatively assess the changes in fungal burden over time, we determined the concentration of galactomannan (GM) in the supernatant of the homogenized fungus ball solution. Consistent with the pathologic findings, GM remained detectable within the fungus balls for more than three months, although the fungal burden gradually decreased (Fig. 3B). It is noteworthy that certain fungus balls disappeared after day 54, whereas in other cases, there was minimal or no reduction in fungal burden (Fig. 3B).

**Figure 3.**
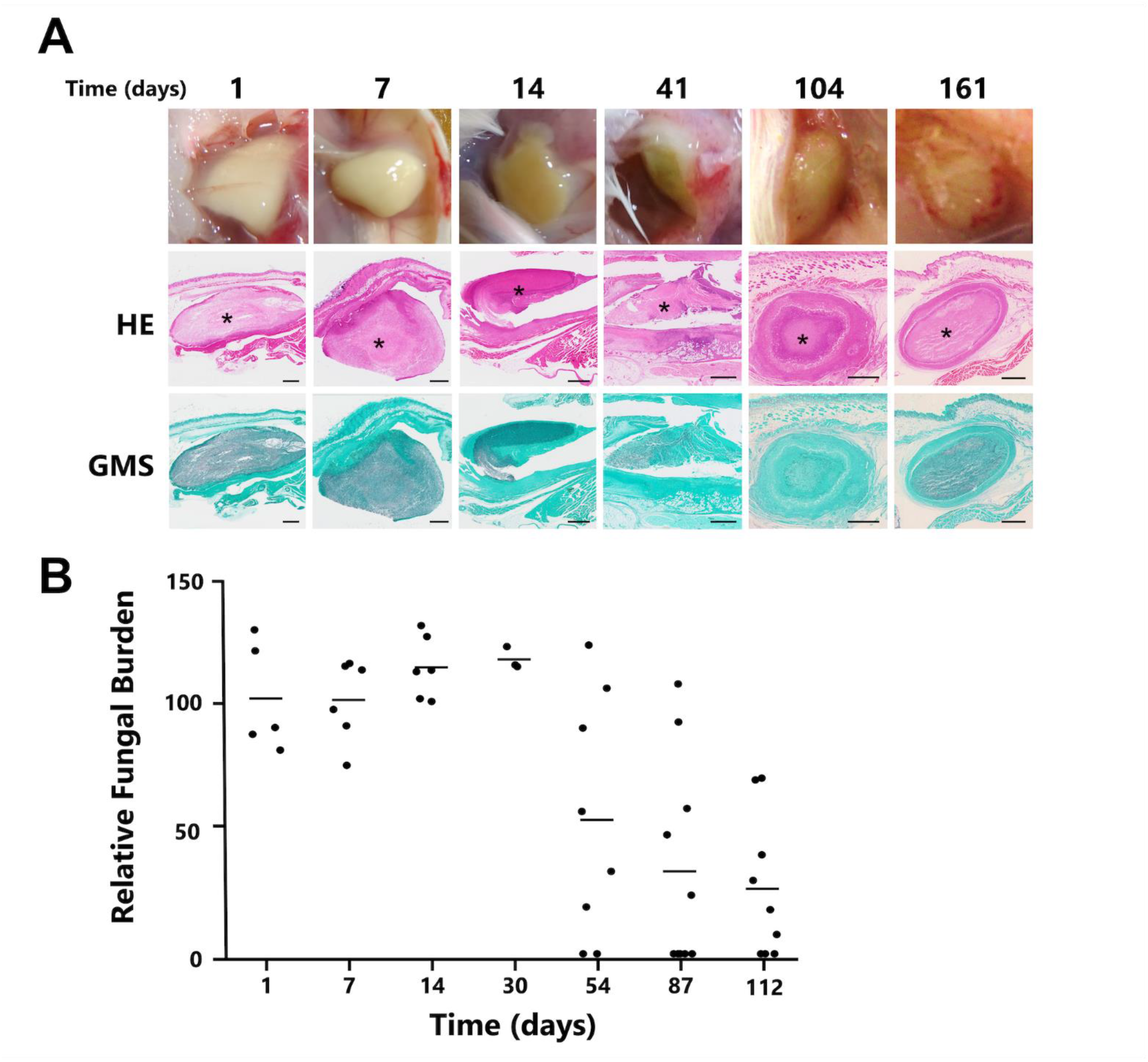
(A) Temporal histopathology of days 1,7, 14, 41, 104, 161 after implantation of dead fungus balls in mice. Visual photographs, H&E staining, and GMS staining for each specimen are shown in order from top to bottom. Asterisks indicate fungus balls. H&E staining shows inflammatory cell infiltration into the fungus ball through day 41 and fibrous structure around the fungus ball after day 104, while GMS staining shows remaining hyphal mass even on day 161. Bars, 1 mm. (B) The time course of fungal burden was measured from homogenized fungus ball solutions using the Bio-Rad Platelia GM assay. If the fungus ball disappeared, it was counted as zero. Scores from each sample are plotted, and the group mean is shown by a bar. The results are represented by 3 to 12 mice per group per time point.

### Differential infiltration of neutrophils and macrophages into the fungus ball

Neutrophils were recruited immediately after implantation and progressively infiltrated into the fungus ball over time (Fig. 4A). Macrophages did not accumulate around the fungus ball until two weeks after implantation and remained on the surface of the fungus ball (Fig. 4A). The accumulation of both neutrophils and macrophages persisted for over three months, with a monthly increase in their area (Fig. 4B, C).

**Figure 4.**
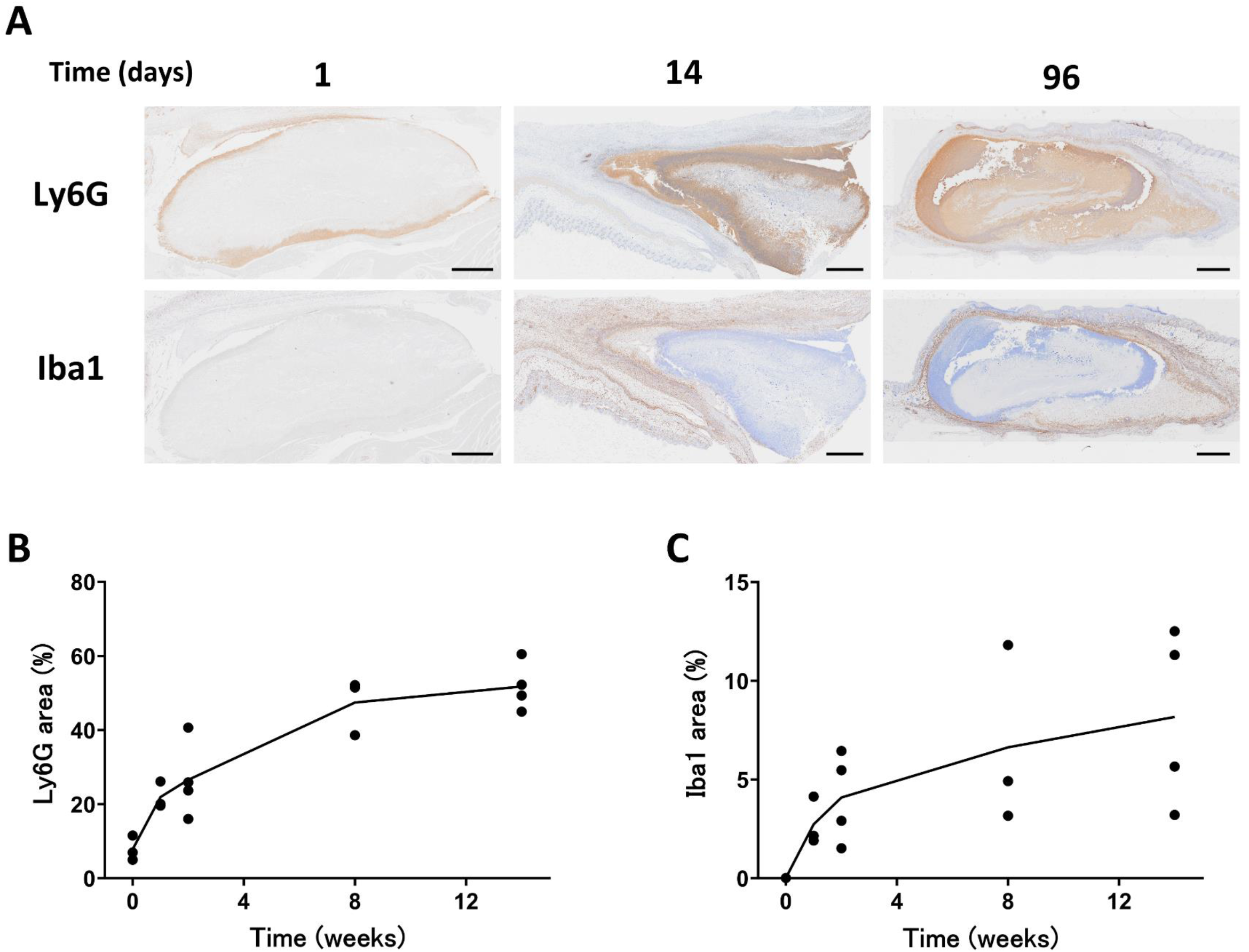
Sections of fungus balls and adjacent tissues were stained for markers associated with (Ly6G) and macrophages (Iba1). (A) Ly6G-positive cells penetrated the fungus balls interior from day 1 following implantation. Conversely, Iba1-positive cells were observed later, surrounding the surface of the fungus balls. The percentage of the area occupied by Ly6G-positive and Iba1- positive cells from each sample are plotted, and increased over time (B, C). Bars, 1 mm.

### Changes in cytokine concentration inside the fungus balls over time

To explore the nature of the inflammatory response to dead fungus balls, we measured the concentrations of eight cytokines from homogenized fungus ball solutions. There was substantial mouse-to-mouse variation in the cytokine levels, which made it difficult to ascertain the profile of cytokine response (Fig. 5). At least IL-1β, TNF-α, IFN-γ, IL-17, and VEGF were elevated for several weeks after fungus ball implantation. IL-10 levels remained below the limit of detection at almost all time points.

**Figure 5.**
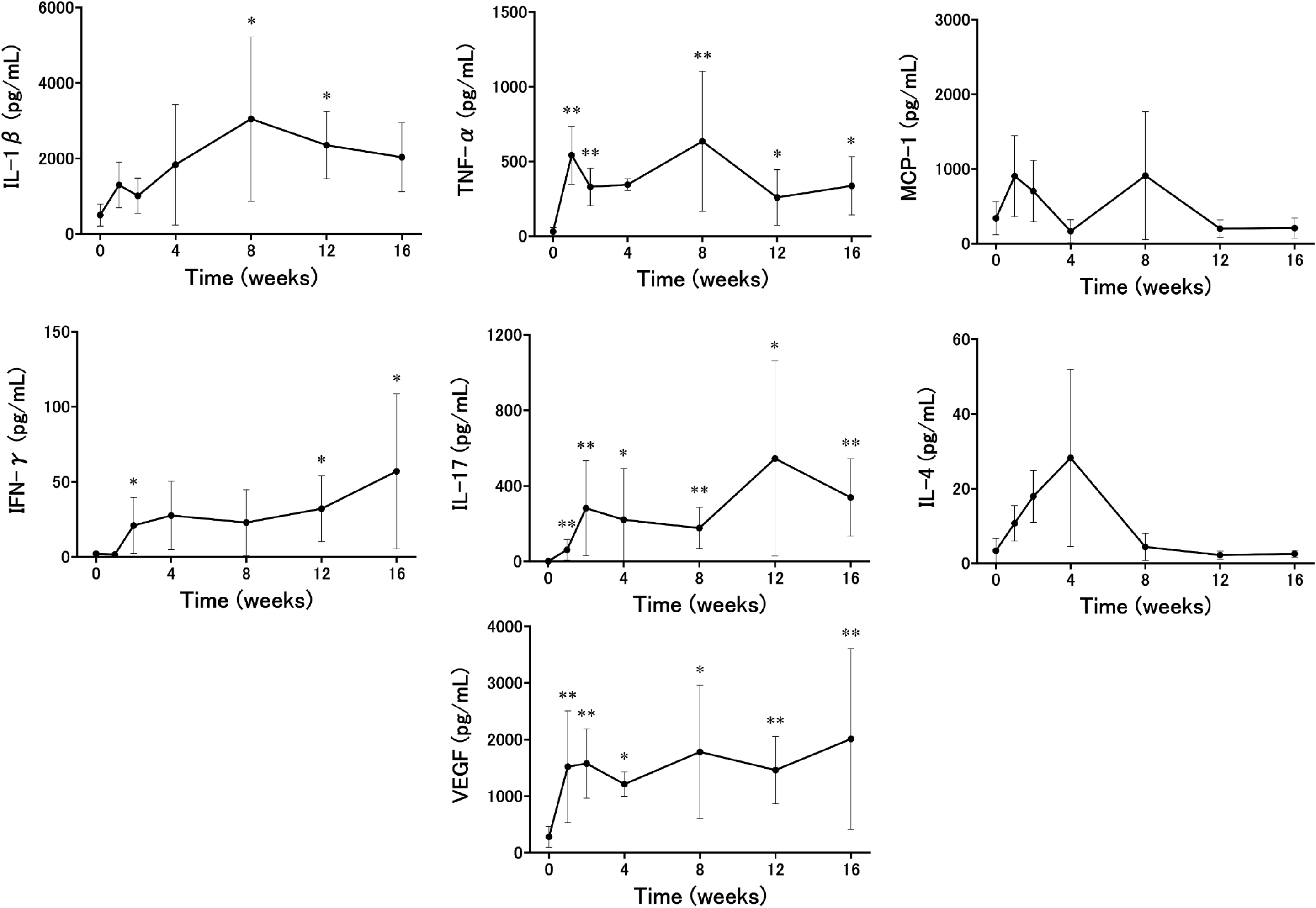
Dead fungus balls induce early proinflammatory mediator release. The concentration of the indicated cytokines/chemokines was measured from homogenized fungus ball solutions 0, 1, 2, 4, 8, 12, and 16 weeks post-implantation using the MILLIPLEX mouse cytokine/chemokine assay. Bars represent the means ± standard errors of the means from at least two independent experiments (total, n = 3 to 6 per group). IL-10 levels are not shown in the figure because all samples were below the detection limit. *P* values were determined by the Mann-Whitney U test compared to baseline (0 week). *, *P* < 0.05; **, *P* < 0.01.

### Characteristics of cells disposing of dead fungus ball

One day after the dead fungus balls were implanted, neutrophils began to accumulate around the killed hyphae; after two weeks, the inflammatory infiltrate around the hyphae shifted from neutrophils to macrophages, and by the third month, macrophages became the primary inflammatory cells on the surface of the fungus balls (Fig. 6A). Interestingly, the macrophages that aggregated around the fungus balls and contained some of the disrupted hyphae swelled over time. These swollen macrophages were positive for Oil red O staining, indicating that they were foam cells containing fat droplets (Fig. 6A). Quantification of the ratio of Oil red O-positive areas to the cross-sectional area of fungus balls showed a significant increase on day 98 compared to day 14, indicating an increase in foam cells (Fig. 6B, p < 0.05). In order to investigate the reasons behind the accumulation of foam cells, characterized by an excess of intracellular lipids, around the fungus ball, we evaluated the expression of peroxisome proliferator-activated receptor gamma (PPAR-γ), a critical transcription factor in lipid metabolism (Fig. 7). While some reports suggest that inflammation-induced reduction in PPAR-γ expression promotes foam cell formation in macrophages [25], we observed PPAR-γ positivity in the foam cells surrounding the fungus ball. To further elucidate the origin of macrophages around the fungus ball, we compared the expression of cell markers CD163 and CD206 with those of skin-resident macrophages (Fig. 7). Skin-resident macrophages displayed positivity for both CD163 and CD206, whereas macrophages around the fungus ball tested negative for both markers. These findings suggest that macrophages around the fungus ball exhibit distinct characteristics from skin-resident macrophages, possibly originating from other locations such as the bone marrow.

**Figure 6.**
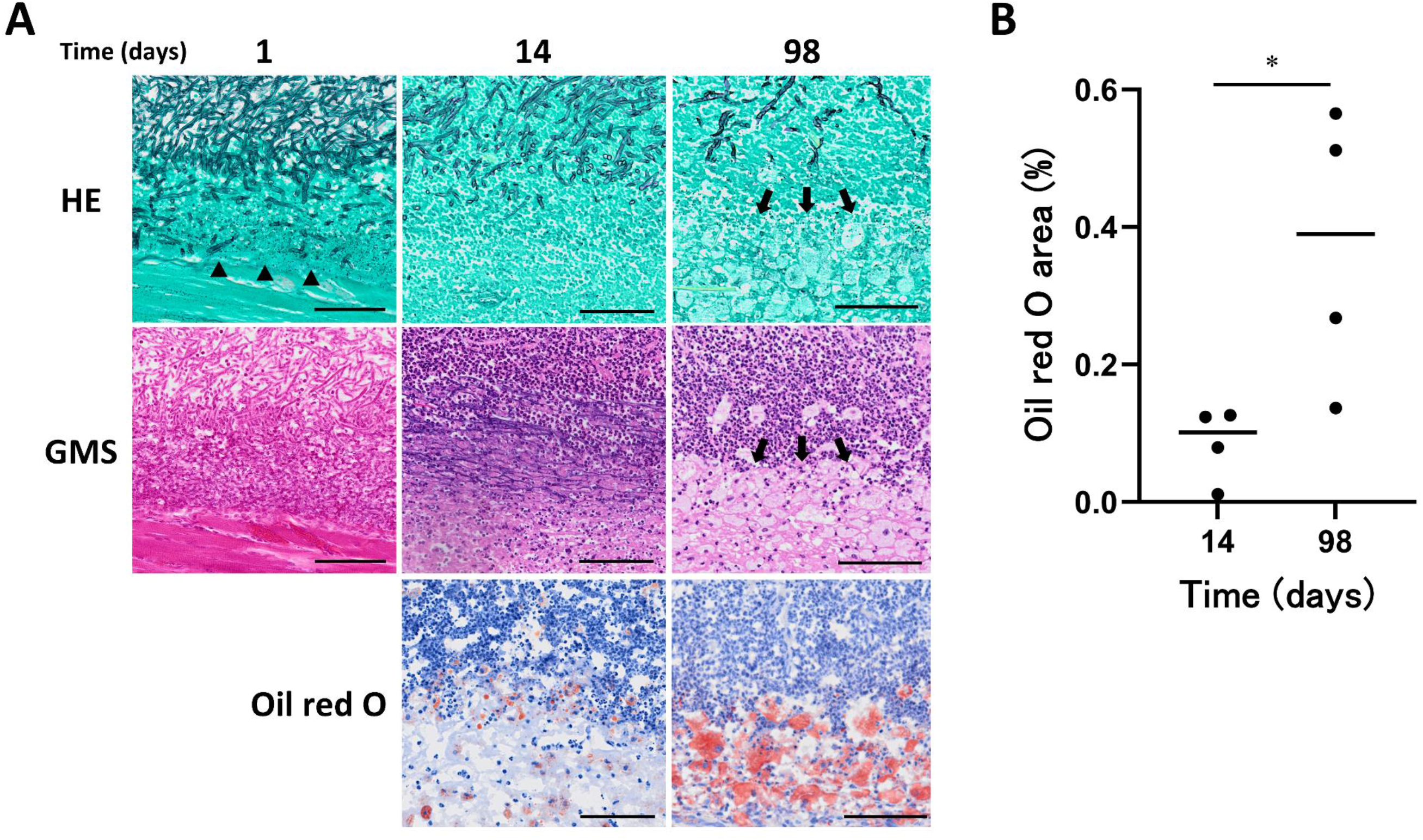
(A) Detailed histopathology of the surface of the fungus ball on days 1, 14, and 98. H&E staining shows the inflammatory cell infiltrate shifting from neutrophils to macrophages. GMS staining of the same area shows that the hyphae were broken into pieces by neutrophils on day 1 and phagocytosed by macrophages on day 98. Black arrowheads and arrows indicate fragmented hyphae and swollen macrophages phagocytosing them, respectively. The Oil Red O staining shows that the macrophages have transformed into foam cells containing fat droplets through day 98. Bars. 100 μm. (B) The percentage of area occupied by Oil red O-positive cells in the fungus ball and surrounding tissue was significantly increased on day 98 compared to day 14. Bars represent the means ± standard errors of the means from at least two independent experiments (total, n = 4 per group). *P* values were determined by the Mann-Whitney U test. *, *P* < 0.05.

**Figure 7.**
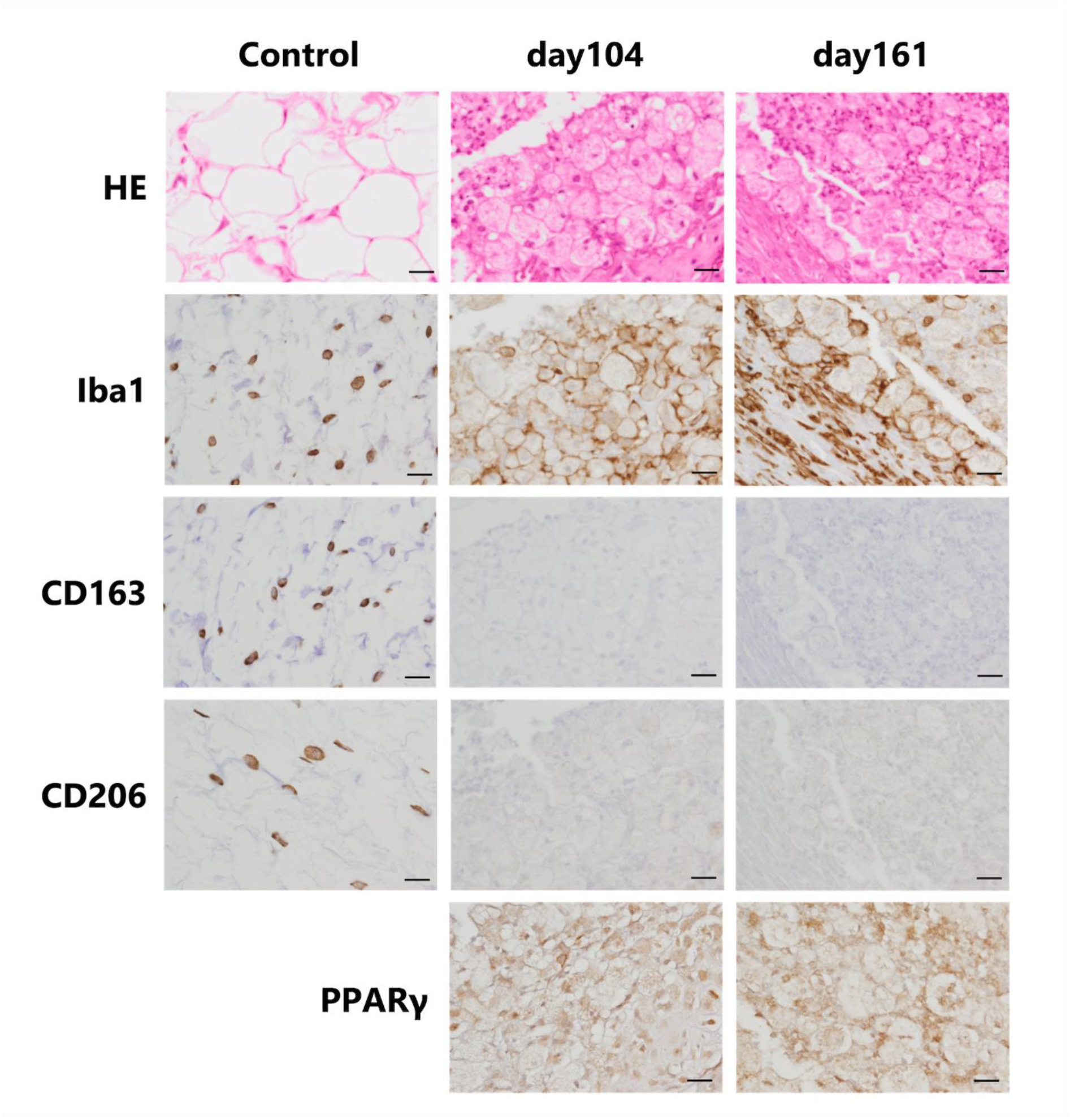
Control sections of uninfected skin and sections from the surface of fungus balls on days 104 and 161 were subjected to immunohistochemistry (IHC) to label macrophage-related (Iba1) and macrophage activation (CD163 and CD206) markers. Iba1-positive cells were detected in the skin, co-expressing CD163 and CD206. In contrast, many Iba1-positive cells were found on the surface of the fungus ball on days 104 and 161, but the stains for CD163 and CD206 were negative. The fungus ball surface was also immunostained for peroxisome proliferator-activated receptor gamma (PPAR-γ) protein, a lipid metabolism regulator. The macrophages surrounding the fungus ball on days 104 and 161 were positive for PPAR-γ. Bars, 20 μm.

### Macrophage damage and morphological changes caused by dead hyphae

We investigated the interaction between dead hyphae and macrophages in vitro. We incubated bone marrow derived macrophages (BMDMs) with killed *A. fumigatus* hyphae and found that there was significant cytotoxicity (Fig. 8A). Maximal cytotoxicity was achieved by direct contact between the hyphae and the BMDMs. When the killed hyphae were separated from the macrophages by filter inserts, the cytotoxic effect was significantly reduced (Fig. 8A). We then investigated whether the formation of foam cells observed in the mouse model could be recapitulated in vitro. We cultured RAW264.7 cells with dead hyphae and stained them with Oil-red-O, which revealed a high level of lipid accumulation, similar to cells that had engulfed cholesterol (Fig. 8B). Our results suggest that phagocytosis of killed hyphae induces macrophages to accumulate lipids and transform into foam cells.

**Figure 8.**
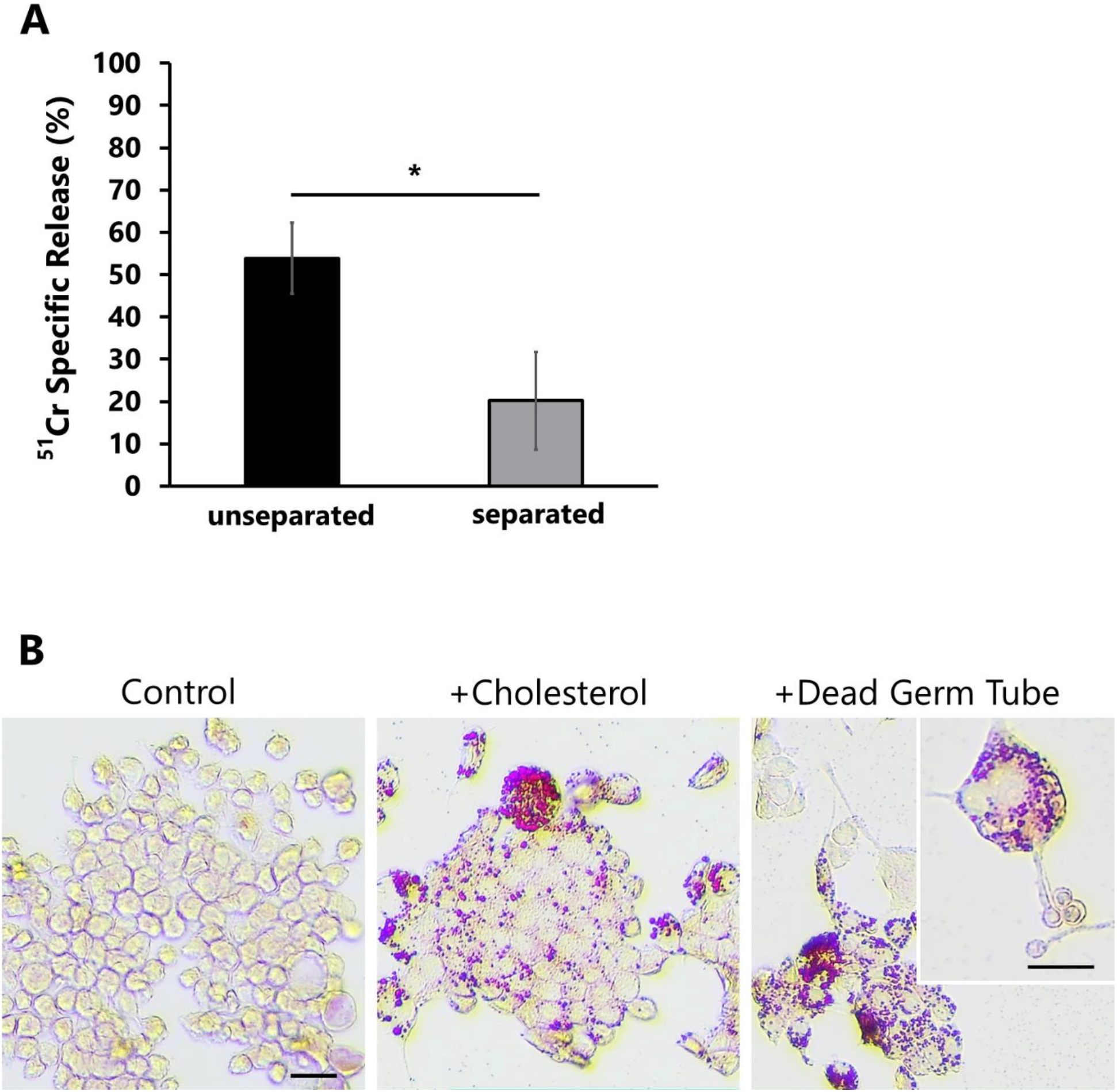
Impact of macrophages contact with heat-killed *A. fumigatus* germ tubes. (A) ^51^Cr-release assay of macrophages and germ tubes with and without transwell separation. BMDM and germ tubes were plated with and without separation for 48 hrs. The results are the mean ±SD of 3 experiments, each performed in triplicate. * Statistically significant, p < 0.01. (B) Oil Red O staining of RAW264.7 cells incubated with either cholesterol or dead *A. fumigatus* germ tubes. Control, the same staining was performed on RAW 264.7 cells alone. Many RAW264.7 cells transformed into foam cells after phagocytosing dead germ tubes. Bars, 20 μm.

## DISCUSSION

Research on the chronic interactions between the fungus and host during aspergilloma formation has been limited due to the lack of an appropriate model in which fungus balls can persist in immunocompetent mice [17]. Our animal model, which involves implanting a fungus ball into an air-filled subcutaneous cavity in a mouse, provided pathological findings that closely resemble those found in clinical aspergillomas. We succeeded in reproducing key features of human aspergilloma by implanting only a dead fungal ball without live fungi. Over the 3-month observation period, neutrophils infiltrated the interior of the fungus ball, while macrophages surrounded its surface. Additionally, macrophages in direct contact with dead hyphae on the surface of the fungus ball were transformed into foam cells. These findings suggest that inability of macrophages to digest dead hyphae causes them to become dysfunctional and transform into foam cells. The chronic immune response observed in this model holds promise to provide valuable insights into the formerly mysterious interaction between host and aspergilloma.

Histopathological analysis of this model suggests that the primary components of clinical aspergilloma are dead fungal hyphae. Transplanting live fungus balls into subcutaneous cavities of healthy mice resulted in the invasion of fungal hyphae into the surrounding tissues. This finding suggests that significant exposure to live *A. fumigatus* hyphae can induce tissue invasion even in healthy hosts. Conversely, transplantation of heat-killed fungus balls closely replicated the histopathological findings observed in human aspergillomas. In aspergillomas in humans, *Aspergillus* may undergo repeated cycles of growth and decay within the cavity, ultimately leading to the formation of aspergilloma, characterized by a dead organism at its core [26].

Importantly, in our model, the aspergilloma persisted within the cavity for over three months, even though it consisted entirely of dead fungal hyphae. The fungal load declined after day 30 post-implantation. By day 112, while the fungus ball disappeared in some mice, residual fungal elements remained present in a substantial majority of the animals.

A notable histopathological finding in this model was that the host response to the dead fungus ball resulted in the generation of fibrotic cavity walls accompanied by abundant neovascularization, similar to a human aspergilloma [15]. The results of the persistently elevated VEGF concentration inside the aspergilloma of this animal model suggest that the placement of the dead fungus ball stimulated angiogenesis around the cavity wall. Indeed, increased expression of VEGF has been reported in clinical cases of aspergillomas [27]. Our results indicate that even if antifungal agents kill all the *Aspergillus* inside the aspergilloma, any remaining dead fungi may cause hemoptysis-causing angiogenesis. Thus, treatment of aspergillomas should focus not only on killing live *Aspergillus*, but also on eliminating the dead organisms.

Our animal model revealed characteristics of a chronic immune response to aspergilloma composed of dead hyphae. Immunohistochemical observations demonstrated early infiltration of neutrophils into the fungus ball, followed by macrophages accumulating around the periphery.

While this study did not provide a detailed profile of cytokines within the fungus ball, the concentrations of key inflammatory cytokines, including IL-1β, TNF-α, IFN-γ, and IL-17, were observed to increase within the fungus ball following transplantation. These results suggested activation of the innate immune response, the Th1 response, and the Th17 response. Studies on nasal tissues obtained during surgery for sinus fungus balls have reported significant infiltration of neutrophils and macrophages and increased IL-1β levels [28]. Furthermore, it has been reported that patients with severe CPA also exhibit elevated serum levels of IL-1β [29]. It has also been reported that serum IL-1β, TNF-α, and IFN-γ levels are significantly increased in CPA patients compared to healthy controls, while IL-4 and IL-10 levels are not [30]. These findings align with the results observed in this mouse model. An important point is that our animal model enables us to analyze the immune response over 16 weeks. This prolonged duration is crucial for understanding the evolution of the host inflammatory response to the fungus ball.

Our model showed that neutrophils were recruited within the fungus ball immediately after implantation, while in the chronic phase, macrophages surrounded the fungus ball. The presence of fungal hyphal fragments within macrophages suggested an unsuccessful attempt to phagocytize and process the fungal debris. Furthermore, the transformation of macrophages into foam cells upon phagocytosis of fungal hyphal fragments implies that the macrophages became functionally impaired [31]. As indicated by the immunohistochemistry (IHC) results, the resulting foam cells were demonstrated to be M1-like macrophages that were likely recruited bone marrow cells rather than resident macrophages from the skin [32]. PPAR-γ, a protein associated with lipid uptake and regulation of lipid metabolism, is suggested to play a role in the formation of foam cells. We hypothesize that they formed with macrophages exceeded their capacity to incorporating lipids contained within the dead hyphae [33].

The observed foam cells in this study, known as therapeutic targets in atherosclerosis, are also associated with chronic inflammation caused by various infections and metabolic disorders [31]. Foam cells form when the lipid content of macrophages exceeds their ability to maintain lipid homeostasis, compromising the critical immune function of macrophages and leading to chronic inflammation [31]. The process of foam cell formation varies based on the disease. In atherosclerosis, it occurs due to the build-up of cholesterol within macrophages, whereas in tuberculosis, it is caused by the accumulation of triglycerides [34,35]. While studies on fungal infections that demonstrate the presence of foam cells are limited, a study of *Histoplasma capsulatum* infection has shown that β-glucan in the cell wall triggers the formation of lipid bodies in leukocytes through CD18, Dectin-1, and TLR2-dependent mechanisms [36].

Additionally, the phagocytosis of heat-killed *Candida albicans* by macrophages has been shown to induce foam cell formation and inflammation through upregulation of FABP4 [37]. Although there are few reports of foam cell involvement in aspergillosis, foam cells have been observed in orbital aspergillosis where fine needle aspiration biopsy was performed [38]. In a series of 16 cases of aspergillosis, 75% of the cases showed a tissue reaction characterized by foreign body giant cells with foam cells [39].

The current study revealed that the long-term interaction of the host with dead *Aspergillus* hyphae results in foam cell formation, suggesting a potential link to chronic inflammation. Furthermore, in vitro experiments demonstrated that direct phagocytosis of killed hyphae resulted in significant damage to macrophages, suggesting that the interaction of macrophages with components of *A. fumigatus*, such as the cell walls, cell membranes, or secondary metabolites, may be involved in foam cell conversion. Further investigation of the mechanism of foam cell transformation may lead to a better understanding of the causes of chronic inflammation and macrophage dysfunction in aspergilloma.

This aspergilloma model, in which *A. fumigatus* fungus balls are implanted in the subcutaneous cavity, provides essential insights into chronic host-aspergilloma interactions. However, there are notable differences between this mouse model and the human disease. This model allows the analysis of processes that occur after the formation of an aspergilloma but not during its formation. Furthermore, while human aspergillomas often have mobile fungus balls within the cavity [26], the fungus ball in this mouse model is primarily adherent to the tissue. Consequently, the prolonged contact between the fungus ball and host tissue in this model may accelerate the host response. Nevertheless, it is worth noting that spontaneous reduction or resolution of size has been reported in around 10% of clinical cases of aspergilloma without specific treatment [41]. This suggests the possibility that a similar immune response, as observed in this mouse model, might occur in some aspergilloma patients. Finally, it is important to mention that while the fungus balls in this study were composed entirely of dead organisms, in patients with aspergillomas, it is typical for living hyphae to be present on the surface [11]. At present, our team is examining an animal model of aspergilloma that includes living hyphae on the surface while preserving a dead organism’s core.

In conclusion, using this novel mouse model, we have characterized the critical events that occur during the interaction of an *A. fumigatus* fungus ball in a cavity with the host. The results suggest that macrophages play an important role in the chronic phase of the immune response to aspergilloma and that the conversion of macrophages into foam cells during phagocytosis of dead hyphae may lead to prolongation of the aspergilloma. Understanding how the host effectively processes dead *Aspergillus* hyphae could potentially lead to the development of novel therapeutic approaches for aspergilloma. Because of its minimally invasive and simple nature, we believe that this mouse model can be applied to a wider range of experiments, potentially leading further insights in the future.

## METHODS

### Mice

Female ICR mice aged 6 to 9 weeks were purchased from Japan SLC, Inc. (Shizuoka, Japan). The animals were kept under standardized, sterile environmental conditions (room temperature 24°C, relative humidity 50%) on a 12-h light-dark cycle and received food and water ad libitum. All the animal experiments were performed in accordance with the recommendations in the Fundamental Guidelines for Proper Conduct of Animal Experiment and Related Activities in Academic Research Institutions under the jurisdiction of the Ministry of Education, Culture, Sports, Science, and Technology. All experimental procedures were approved by the Institutional Animal Care and Use Committee of Nagasaki University (approval number 2012091679).

### Preparation of fungus balls

We used *A. fumigatus* MF367, a clinical isolate obtained from a patient with chronic pulmonary aspergillosis, to establish an aspergilloma mouse model [42]. This strain was cultured on potato dextrose agar at 30°C for 4 to 5 days; conidia were collected by rinsing the culture plates with phosphate-buffered saline containing 1% tween 20 (FUJIFILM Wako Pure Chemical, Tokyo, Japan). The conidial suspension was diluted to a final concentration of 1 × 10^5^ cells/ml. Two hundred microliters of the conidial suspension were added to 20 ml potato dextrose broth and incubated on a rotary shaker at 250 rpm for 48 hours at 30°C. As a result, a large number of fungus balls, with diameters ranging from 3 to 7 mm, were generated. Fungus balls of approximately 5 mm diameter were selected from the medium to minimize variation. Individual fungus balls were selected and transferred to fresh 20 ml potato dextrose broth and incubated on a rotary shaker at 30°C for 24 hours to obtain larger fungus balls, approximately 10 mm in diameter. If dead fungus balls were required, the balls were autoclaved at 121°C for 15 minutes to kill the organisms. Loss of viability of the autoclaved fungus balls was confirmed by their inability to grow in culture. The balls were washed twice with 20 ml of 0.9% saline prior to implantation into mice.

### Fungus ball implantation into mice

On day 0, the mice were anesthetized with an intraperitoneal injection of medetomidine (Kyoritsu Seiyaku Corporation, Tokyo, Japan; 0.3 mg/kg), midazolam (Sandoz, Holzkirchen, Germany; 4.0 mg/kg), and butorphanol (Meiji Seika Pharma Co. Ltd., Tokyo, Japan; 5.0 mg/kg). Subsequently, 10 ml of room air was injected under the skin on the back of the mouse to create an air-filled cavity, into which a fungus ball was inserted (Fig. 1). The cavity was maintained by periodic injection of air 2-3 times per week.

### Immunization with *A. fumigatus* hyphae prior to fungus ball implantation

A 4-5 mm live fungus ball was homogenized in a 2 ml sealed vial containing a 6 mm stainless steel beads and 1 ml saline using a BMS-M10N21 homogenizer (BMS, Tokyo, Japan) at 1500 rpm for 1 minute. Before fungus ball implantation, mice were injected intraperitoneally with 1 ml of a 10-fold dilution of the homogenized solution twice weekly for 4 to 8 weeks.

Homogenates prepared by this method contain live fungus. The immunization status of the mice against *Aspergillus* was confirmed by measuring the antibody levels in the sera of the immunized mice by immunoprecipitation using the FSK1 *Aspergillus* immunodiffusion system (Microgen Bioproducts Ltd., United Kingdom). The test was considered positive if at least one precipitation line was detected by visual observation.

### Histopathological and immunopathological staining

For histopathologic analysis, mice were sacrificed, and the fungus balls and surrounding tissues were dissected out and fixed with 10% formalin. The specimens were embedded in paraffin and sectioned at 3 μm with the fungus balls as the largest section surface. The sections were stained with Grocott’s methenamine silver (GMS), hematoxylin and eosin (H&E). The sections from paraffin-embedded samples were processed for immunohistochemistry by methods described previously [32]. The following antibodies were used to detect neutrophils, macrophages, and peroxisome proliferator-activated receptor gamma (PPAR-γ): anti-Ly6G (rabbit monoclonal, ab238132, Abcam), anti-Iba-1 (rabbit polyclonal, 019-19741, WAKO, Tokyo, Japan), anti- CD163 (rabbit monoclonal, EPR19518, Abcam,), anti-CD206 (rabbit polyclonal, ab64693, Abcam), and anti-PPAR-γ (rabbit polyclonal, ab59256, Abcam), respectively. The samples were incubated with primary antibodies followed by HRP-labeled secondary antibodies (Nichirei, Tokyo, Japan). The reaction was visualized using the Diaminobenzidine (DAB) system (Nichirei). In addition, the lipid content was evaluated by euthanizing the mice, surgically removing the fungus balls and adjacent tissues, and preparing frozen sections. The lipid in the frozen sections were visualized using the Oil Red O staining protocol. The sections were gently rinsed with distilled water and then immersed in 60% isopropanol for 20-30 seconds. They were stained with Oil Red O for 60 minutes, followed by washing with 60% isopropanol and distilled water. The nuclei were stained with Mayer’s hematoxylin, rinsed with water for 10 minutes, and the slides were coverslipped.

### Image analysis

GMS, H&E, Oil red O, and immunohistochemical staining results were observed using OlyVIA software (Olympus Olyvia 3.4) after scanning with a slide scanner (VS200, Olympus). Image analysis was performed using the Fiji software (Image J version 2.9) to determine the number of neutrophils, macrophages, and cells containing lipid droplets inside the fungus balls. Using ImageJ, the percent coverage of DAB and Oil red O staining in the fungus ball region was quantified and evaluated over time. First, to evaluate leukocytes clustered inside and around the fungus ball, a 500 μm margin was set from the fungus ball outline using a freehand selection tool (Figure S3A). We used the Clear Outside tool to erase areas outside the margin and removed the counterstained nuclei from the image using the Threshold Colour plugin to solely assess the positive DAB or Oil red O staining (Fig. S3B). A threshold was set to ensure analysis of the cell population of interest that best captured positive staining while minimizing background. Using the threshold, we defined positive immunohistochemistry in red and calculated the area (Fig. S3C). We also set a different threshold that gives the total area of the fungus ball and the margin (Fig. S3D). The percentages of DAB and Oil red O positive areas were calculated by dividing the respective positive area by the total area and multiplying by 100.

### Galactomannan assay

Fungus balls, collected on days 1 to 112 after implantation, were homogenized with a 6 mm stainless steel bead, 200 mg of 0.6 mm zirconia/silica beads, and 1 ml saline using the BMS- M10N21 homogenizer at 1500 rpm for 5 minutes. After clarifying the homogenate by centrifugation, the supernatant was collected. The fungal burden of each fungus ball was determined by measuring the galactomannan (GM) content in the supernatant [43] using the Platelia *Aspergillus* enzyme immunoassay kit (Bio-Rad, Hercules, CA) according to the manufacturer’s instructions.

### Cytokine analysis

The fungus ball supernatant was collected as described above. The IL-1ꞵ, IL-4, IL-10, IL-17, IFN-γ, TNF-α, MCP-1, and VEGF content was quantified using a custom-made Milliplex Mouse Cytokine/Chemokine Panel (Merck Millipore, USA), a magnetic bead-based multiplex immunoassay following the manufacturer instructions. Fluorescence was quantified using a Luminex 200 instrument (Luminex Corporation, USA), and data were analyzed with MILLIPLEX Analyst software version 5.1. Cytokine levels in the supernatant were corrected for fungal burden obtained by GM assay.

### In vitro cytotoxic analysis of *A. fumigatus* dead hyphae on macrophages

Bone marrow-derived macrophages (BMDMs) were isolated from 6-week-old mice (Taconic Laboratories). The cells were differentiated into macrophages with 50 ng/ml macrophage colony–stimulating factor (#576404, BioLegend) and maintained by incubation in Dulbecco’s Modified Eagle’s Medium (DMEM) (American Type Culture Collection) with 10% fetal bovine serum (FBS) (Gemini Bio-Products), 1% streptomycin and penicillin within ten days before use. To determine the extent of the host cell damage caused by dead *A. fumigatus* hyphae, a ^51^Cr release assay was used as previously described [44]. The hyphae were generated by incubating *A. fumigatus* conidia in Sabouraud broth at 37°C for 8 h and then killed by heating at 95°C for 10 minutes. The killed hyphae were rinsed extensively in phosphate-buffered saline prior to use in the experiments. The day before the experiment, 2.5 × 10^5^ cells per well of BMDM were labeled with ^51^Cr (ICN Biomedicals) overnight in a 24-well tissue culture plate. The following day, the cells were rinsed twice with Hanks’ Balanced Salt Solution to remove the unincorporated ^51^Cr, then 7.5 × 10^5^ hyphae in 1 ml of tissue culture medium were added to each well. After 48 h of incubation, 0.5 ml of the medium above the cells were collected, and the cells were lysed with 6 N NaOH and rinsed twice with RadiacWash (Biodex Medical Systems). Both the 0.5 ml medium collection and the combined collection of lysed cells were measured by a gamma counter. The spontaneous release of ^51^Cr was determined using uninfected BMDMs processed in parallel. The specific release of ^51^Cr was calculated using our previously described formula [45]. To evaluate the effect of direct contact on BMDM damage, the killed hyphae were added to transwells with 3 µm pores (Transwell Permeable Supports, Corning Inc.) that were suspended above the BMDMs. Each experiment was performed in triplicate and repeated three different times.

### In vitro lipid droplets staining

RAW264.7 cells (a mouse peritoneal macrophage cell line) were cultured in DMEM supplemented with 10% heat-inactivated FBS in humidified 5% CO_2_ at 37°C. The cells were plated (2.5 × 10^4^ cells per well in 24-well plates), cultured in DMEM with 10% FBS for 20 h, and incubated with 1.0 × 10^5^ killed hyphae for an additional 72 h. As a positive control, 10 μM cholesterol was added to RAW264.7 cells and cultured for 72 h. The cells were stained using the Oil-red-O stain kit (Bio Mirai Koubou, Tokyo, Japan) according to the manufacturer’s instructions.

### Statistical analyses

Statistical significance was determined using the Mann-Whitney U test. *P*-value < .05 was considered statistically significant. The statistical analyses were performed using JMP Pro 17 software (SAS Institute, Cary, NC, USA).

## ACKNOWLEDGMENTS

We thank Yuichiro Hashimoto, Naoki Ito, Ryo Urano, and Minori Kikuchi for the creation of the aspergilloma mouse model.

## CONTRIBUTORS

The author’s contribution is as follows: Conceptualization, MT, KI, and RH; methodology, MT, YN, HL, KI, and RH; validation, YN, MT, RH, HN, YI, TH, K Takeda, and NI; formal analysis, RH; investigation, YN, MT, KN, HL, YK, and RH; data curation, RH; writing—original draft preparation, RH; writing—review and editing, RH, MT, KN, HL, YK, SGF and KI; visualization, RH; supervision, T Takazono, T Tanaka, AW, AF, KY, HM, SGF, K Takayama, and KI; project administration, MT; funding acquisition, MT, KN, AW, and SGF; All authors have read and agreed to the published version of the manuscript.

## REFERENCES

1. Denning DW, Chakrabarti A. Pulmonary and sinus fungal diseases in non-immunocompromised patients. Lancet Infect Dis. 2017;17: e357–e366.

2. Smith NL, Denning DW. Underlying conditions in chronic pulmonary aspergillosis including simple aspergilloma. Eur Respir J. 2011;37: 865–872.

3. Lee JG, Lee CY, Park IK, Kim DJ, Chang J, Kim SK, et al. Pulmonary aspergilloma: Analysis of prognosis in relation to symptoms and treatment. J Thorac Cardiovasc Surg. 2009;138: 820–825.

4. Singh V. Fungal Rhinosinusitis: Unravelling the Disease Spectrum. J Maxillofac Oral Surg. 2019;18: 164–179.

5. Garvey J, Crastnopol P, Weisz D, Khan F. The surgical treatment of pulmonary aspergillomas. J Thorac Cardiovasc Surg. 1977;74: 542–547.

6. Citak N, Sayar A, Metin M, Pekçolaklar A, Kök A, Akanıl Fener N, et al. [Results of surgical treatment for pulmonary aspergilloma with 26 cases in six years: a single center experience]. Tuberk Toraks. 2011;59: 62–69.

7. Denning DW, Pleuvry A, Cole DC. Global burden of chronic pulmonary aspergillosis as a sequel to pulmonary tuberculosis. Bull World Health Organ. 2011;89: 864–872.

8. Bongomin F, Gago S, Oladele RO, Denning DW. Global and Multi-National Prevalence of Fungal Diseases—Estimate Precision. Journal of Fungi. 2017;3: 57.

9. Denning DW, Cadranel J, Beigelman-Aubry C, Ader F, Chakrabarti A, Blot S, et al. Chronic pulmonary aspergillosis: rationale and clinical guidelines for diagnosis and management. Eur Respir J. 2016;47: 45–68.

10. Kosmidis C, Denning DW. The clinical spectrum of pulmonary aspergillosis. Thorax. 2015;70: 270–277.

11. Corrin B, Nicholson AG. Pathology of the Lungs E-Book: Expert Consult: Online and Print. Elsevier Health Sciences; 2011.

12. Dufour X, Kauffmann-Lacroix C, Ferrie J-C, Goujon J-M, Rodier M-H, Karkas A, et al. Paranasal sinus fungus ball and surgery: a review of 175 cases. Rhinology. 2005;43: 34–39.

13. Nomura K, Asaka D, Nakayama T, Okushi T, Matsuwaki Y, Yoshimura T, et al. Sinus fungus ball in the Japanese population: clinical and imaging characteristics of 104 cases. Int J Otolaryngol. 2013;2013: 731640.

14. Liu X, Liu C, Wei H, He S, Dong S, Zhou B, et al. A retrospective analysis of 1,717 paranasal sinus fungus ball cases from 2008 to 2017. Laryngoscope. 2020;130: 75–79.

15. Tochigi N, Ishiwatari T, Okubo Y, Ando T, Shinozaki M, Aki K, et al. Histological study of chronic pulmonary aspergillosis. Diagn Pathol. 2015;10: 153.

16. van de Veerdonk FL, Gresnigt MS, Romani L, Netea MG, Latgé J-P. Aspergillus fumigatus morphology and dynamic host interactions. Nat Rev Microbiol. 2017;15: 661–674.

17. Arastehfar A, Carvalho A, Houbraken J, Lombardi L, Garcia-Rubio R, Jenks JD, et al. Aspergillus fumigatus and aspergillosis: From basics to clinics. Stud Mycol. 2021;100: 100115.

18. Urb M, Snarr BD, Wojewodka G, Lehoux M, Lee MJ, Ralph B, et al. Evolution of the Immune Response to Chronic Airway Colonization with Aspergillus fumigatus Hyphae. Infect Immun. 2015;83: 3590–3600.

19. Wang F, Zhang C, Jiang Y, Kou C, Kong Q, Long N, et al. Innate and adaptive immune response to chronic pulmonary infection of hyphae of Aspergillus fumigatus in a new murine model. J Med Microbiol. 2017;66: 1400–1408.

20. Sawasaki H, Horie K, Naito Y, Watabe S, Tajima G, Mizutani Y. Experimental pulmonary aspergilloma. Mycopathol Mycol Appl. 1967;32: 265–274.

21. Nakamura J. Experimental Aspergilloma in the Pleural Cavity of the Rabbit. Kawasaki Med J. 1990. Available: https://kwmed.repo.nii.ac.jp/?action=pages_view_main&active_action=repository_view_main_item_detail&item_id=306&item_no=1&page_id=33&block_id=41

22. Yano T. Growth of Experimental Fungus Balls in the Pleural cavity of Rabbits. Kawasaki Med J. 1992. Available: https://core.ac.uk/download/pdf/268405337.pdf

23. Supiyaphun P, Sampatanukul P, Sukumalpaiboon P. Benign Aspergillus colonization (Aspergilloma) in the middle ear. Otolaryngol Head Neck Surg. 2001;125: 281–282.

24. Richardson MD, Page ID. Aspergillus serology: Have we arrived yet? Med Mycol. 2017;55: 48–55.

25. Wouters E, Grajchen E, Jorissen W, Dierckx T, Wetzels S, Loix M, et al. Altered PPARγ Expression Promotes Myelin-Induced Foam Cell Formation in Macrophages in Multiple Sclerosis. Int J Mol Sci. 2020;21. doi:10.3390/ijms21239329

26. Kurashima A. Analysis of the course of non-invasive pulmonary aspergillosis. Jpn J Med Mycol. 1997;38: 167–174.

27. Inoue K, Matsuyama W, Hashiguchi T, Wakimoto J, Hirotsu Y, Kawabata M, et al. Expression of vascular endothelial growth factor in pulmonary aspergilloma. Intern Med. 2001;40: 1195–1199.

28. Jiang R-S, Huang W-C, Liang K-L. Characteristics of Sinus Fungus Ball: A Unique Form of Rhinosinusitis. Clin Med Insights Ear Nose Throat. 2018;11: 1179550618792254.

29. Zhan M, Xu B, Zhao L, Li B, Xu L, Sun Q, et al. The Serum Level of IL-1B Correlates with the Activity of Chronic Pulmonary Aspergillosis. Can Respir J. 2018;2018: 8740491.

30. Ren W, Li H, Guo C, Shang Y, Wang W, Zhang X, et al. Serum Cytokine Biomarkers for Use in Diagnosing Pulmonary Tuberculosis versus Chronic Pulmonary Aspergillosis. Individ Differ Res. 2023;16: 2217–2226.

31. Guerrini V, Gennaro ML. Foam Cells: One Size Doesn’t Fit All. Trends Immunol. 2019;40: 1163–1179.

32. Nakagawa T, Ohnishi K, Kosaki Y, Saito Y, Horlad H, Fujiwara Y, et al. Optimum immunohistochemical procedures for analysis of macrophages in human and mouse formalin fixed paraffin-embedded tissue samples. J Clin Exp Hematop. 2017;57: 31–36.

33. Moore KJ, Rosen ED, Fitzgerald ML, Randow F, Andersson LP, Altshuler D, et al. The role of PPAR-gamma in macrophage differentiation and cholesterol uptake. Nat Med. 2001;7: 41–47.

34. Chistiakov DA, Melnichenko AA, Myasoedova VA, Grechko AV, Orekhov AN. Mechanisms of foam cell formation in atherosclerosis. J Mol Med . 2017;95: 1153–1165.

35. Peyron P, Vaubourgeix J, Poquet Y, Levillain F, Botanch C, Bardou F, et al. Foamy macrophages from tuberculous patients’ granulomas constitute a nutrient-rich reservoir for M. tuberculosis persistence. PLoS Pathog. 2008;4: e1000204.

36. Sorgi CA, Secatto A, Fontanari C. Histoplasma capsulatum cell wall β-glucan induces lipid body formation through CD18, TLR2, and dectin-1 receptors: correlation with leukotriene B4 generation and …. The Journal of. 2009. Available: https://www.jimmunol.org/content/182/7/4025.short

37. Haider M, Al-Rashed F, Albaqsumi Z, Alobaid K, Alqabandi R, Al-Mulla F, et al. Candida albicans Induces Foaming and Inflammation in Macrophages through FABP4: Its Implication for Atherosclerosis. Biomedicines. 2021;9. doi:10.3390/biomedicines9111567

38. K, Amita, Kumari A, S, Vijay Shankar, M, Sanjay. Orbital aspergillosis masquerading as lymphoma diagnosed by cytology. Amita K, Kumari A, Vijay SS, Sanjay M Orbital aspergillosis masquerading as lymphoma diagnosed by cytology – A case report Indian J Pathol Oncol. 2018;6: 148–150.

39. Das R, Dey P, Chakrabarti A, Ray P. Fine-needle aspiration biopsy in fungal infections. Diagn Cytopathol. 1997;16: 31–34.

40. Lee SH, Lee BJ, Jung DY, Kim JH, Sohn DS, Shin JW, et al. Clinical manifestations and treatment outcomes of pulmonary aspergilloma. Korean J Intern Med. 2004;19: 38–42.

41. Lang M, Lang AL, Chauhan N, Gill A. Non-surgical treatment options for pulmonary aspergilloma. Respir Med. 2020;164: 105903.

42. Tashiro M, Izumikawa K, Minematsu A, Hirano K, Iwanaga N, Ide S, et al. Antifungal susceptibilities of Aspergillus fumigatus clinical isolates obtained in Nagasaki, Japan. Antimicrob Agents Chemother. 2012;56: 584–587.

43. Sheppard DC, Marr KA, Fredricks DN, Chiang LY, Doedt T, Filler SG. Comparison of three methodologies for the determination of pulmonary fungal burden in experimental murine aspergillosis. Clin Microbiol Infect. 2006;12: 376–380.

44. Liu H, Xu W, Bruno VM, Phan QT, Solis NV, Woolford CA, et al. Determining Aspergillus fumigatus transcription factor expression and function during invasion of the mammalian lung. PLoS Pathog. 2021;17: e1009235.

45. Pongpom M, Liu H, Xu W, Snarr BD, Sheppard DC, Mitchell AP, et al. Divergent targets of Aspergillus fumigatus AcuK and AcuM transcription factors during growth in vitro versus invasive disease. Infect Immun. 2015;83: 923–933.

